# Insights into the genetic diversity of *Mycobacterium tuberculosis* in Tanzania

**DOI:** 10.1101/441956

**Authors:** Liliana K. Rutaihwa, Mohamed Sasamalo, Aladino Jaleco, Jerry Hella, Ally Kingazi, Lujeko Kamwela, Amri Kingalu, Bryceson Malewo, Raymond Shirima, Anna Doetsch, Julia Feldmann, Miriam Reinhard, Sonia Borrell, Klaus Reither, Basra Doulla, Lukas Fenner, Sebastien Gagneux

## Abstract

**Background:** Human tuberculosis (TB) is caused by seven phylogenetic lineages of the *Mycobacterium tuberculosis* complex (MTBC), Lineage 1–7. Recent advances in rapid genotyping of MTBC based on single nucleotide polymorphisms (SNP), allow for rapid and phylogenetically robust strain classification, paving the way for defining genotype-phenotype relationships in clinical settings. Such studies have revealed that, in addition to host and environmental factors, different strains of the MTBC influence the outcome of TB infection and disease. In Tanzania, such molecular epidemiological studies of TB however are scarce in spite of a high TB burden.

**Methods and Findings:** Here we used a SNP-typing method to genotype a nationwide collection of 2,039 MTBC clinical isolates obtained from new and retreatment TB cases diagnosed in 2012 and 2013. Four lineages, namely Lineage 1–4 were identified. The distribution and frequency of these lineages varied across the regions but overall, Lineage 4 was the most frequent (n=866, 42.5%), followed by Lineage 3 (n=681, 33.4%) and 1 (n=336, 16.5%), with Lineage 2 being the least frequent (n=92, 4.5%). A total of 64 (3.1%) isolates could not be assigned to any lineage. We found Lineage 2 to be associated with female sex (adjusted odds ratio [aOR] 2.25; 95% confidence interval [95% CI] 1.38 – 3.70, p<0.001) and retreatment (aOR 1.78; 95% CI 1.00 – 3.02, p=0.040). We found no associations between MTBC lineage and patient age or HIV status. Our sublineage typing based on spacer oligotyping revealed the presence of mainly EAI, CAS and LAM families. Finally, we detected low levels of multidrug resistant isolates among a subset of retreatment cases

**Conclusions:** This study provides novel insights into the influence of pathogen-related factors on the TB epidemic in Tanzania.

## Introduction

Tuberculosis (TB) is the leading cause of mortality due to an infectious disease [1]. In 2017, an estimated 10.0 million people developed TB globally, with 1.3 million dying of the disease. More than 80% of the TB burden lies in the thirty high burden countries [1]. Tanzania is among these countries, with a national average TB notification rate of 129 cases per 100,000; however, some regions show higher notification rates [2]. Like in most sub- Saharan African countries, the HIV epidemic contributes to the high TB incidence in Tanzania, where a-third of the TB patients are co-infected with HIV [2]. Contrarily, drug resistant-TB is still low in this setting [3]. Other risk factors such poverty also influence the epidemiology of TB in Tanzania [4].

Transmission of TB occurs via infectious aerosols, and upon exposure individuals can either develop active disease or remain latent infected [5]. It is estimated that a-quarter of the world′s population is latently TB infected [6], with a 5 – 10% life time risk to develop active TB disease; this risk is 50% in case of HIV co-infected individuals [7].

The complex dynamics of TB infection and disease are determined by the environment, the host and the pathogen [8]. Seven main phylogenetic lineages of the *Mycobacterium tuberculosis* complex (MTBC) lineages (Lineage 1–7) are known to cause TB in humans [9]. These lineages are phylogeographically distributed, reflecting human migration histories and possibly adaptation to different human populations [10–12]. Some genomic differences among the MTBC strains translate into relevant biological and epidemiological phenotypes [13]. In general, strains of the globally distributed lineages, Lineage 2 and 4 or “generalists”, appear to be more virulent in average than those of the geographically restricted lineages, Lineage 5 and 6 or “specialists” [9,13]. Epidemiologically speaking, these phenotypes are demonstrated by indicators such as transmission potential, disease severity and rate of progression from infection to disease [14–17].

Studying genotype-phenotype relationships requires understanding the genetic diversity of MTBC clinical strains in a given clinical setting. In Tanzania few studies have described the genetic diversity of MTBC [18‒21]. These previous work revealed the presence of mainly three lineages; Lineage 1, 3 and 4, which include the EAI, CAS and LAM spoligo families, respectively. Lineage 2, which includes the Beijing family, has only been reported at the lowest frequencies. These previous studies are limited as they only focused on few geographical locations and used spacer oligonucleotide typing (spoligotyping) technique which has limitations for phylogenetically robust strain classification [22,23]. Only one study profiled MTBC on a countrywide scale albeit with low sampling coverage [20].

In this study we used for the first time a robust single nucleotide polymorphism (SNP) typing method to classify the largest so far nationwide collection of MTBC clinical isolates from Tanzania. We then looked for potential associations between the MTBC lineages and the clinical and epidemiological characteristics of the patients.

## Material and Methods

### Study setting

Our study was based on a nationwide convenience sample of sputum smear positive new and retreatment TB cases diagnosed between 2012 and 2013 in Tanzania. The collection was obtained via a platform established for routine TB drug resistance surveillance by the National Tuberculosis Leprosy Program (NTLP) of Tanzania, covering health facilities in all geographical regions of the country. Briefly, smear positive sputa specimens from approximately 25% of new TB cases (obtained by allocating four months of sample collection to each region annually) and from all retreatment cases were sent to zonal reference laboratories (i.e. Central Tuberculosis Reference Laboratory [CTRL] in Dar es Salaam, Bugando Medical Center [BMC] in Mwanza and Kilimanjaro Christian Medical Center [KCMC] in Kilimanjaro), which serve the respective nearby regions for culture. Isolates from the two zonal laboratories, BMC and KCMC were then sent to the CTRL for drug susceptibility testing (DST). For this study we used archived isolates obtained from the CTRL.

### Study population and data collection

We included a total of 2,039 unique (single patient) culture-confirmed TB cases diagnosed between 2012 and 2013, each of whom we could retrieve the respective culture isolate. This study population represents 41% of all culture positive sputum samples processed and 1.6% of all TB notified cases in the country during the study period (S1 Fig). We also obtained corresponding socio-demographic and clinical information collected during patients’ consultation at the respective health facilities. The demographic data collected included age, sex and geographical location of the patients, whereas clinical data included HIV status and disease category (i.e., new case and retreatment case).

### Processing of culture isolates

The smear positive sputa samples were cultured on Löwenstein Jensen (LJ) growth medium according to laboratory protocols. For this study, we included *M. tuberculosis* clinical isolates retrieved from archived LJ media. We then prepared heat inactivated samples for the retrieved clinical isolates by suspending *M. tuberculosis* colonies into 1ml sterile water and heat inactivate at 95°C for one hour.

### Molecular genotyping

We then classified the *M. tuberculosis* clinical isolates into main phylogenetic lineages by TaqMan real-time PCR according to standard protocols (Applied Biosystems, Carlsbad, USA) and as previously described [24]. We also performed 43-spacer spoligotyping on a membrane for a subset of representative *M. tuberculosis* clinical strains following standard protocols [25]. The clinical strains were assigned to spoligotype families using the online database SITVITWEB [26].

### Drug Resistance Genotyping

We selected a subset of clinical isolates from retreatment cases to perform molecular drug resistance testing. We used a previously described multiplex polymerase chain reaction (PCR) to target the hotspot region of *rpoB* gene that confers resistance to rifampicin [27]. The PCR assay targets both the tuberculous and non-tuberculous *Mycobacteria* (MTBC and NTMs, respectively) *rpoB* gene, so we could also rule out the presence of non-tuberculous isolates in our study sample using the assay. The amplified *rpoB* gene product was confirmed by electrophoresis on a 2% agarose gel and sent for Sanger sequencing. We analyzed the resulting sequences by Staden software package [28] and using *M. tuberculosis* H37Rv *rpoB* gene as reference sequence.

### Statistical Analyses

For statistical analyses we applied descriptive statistics to delineate patients′ characteristics. We used Chi-square or Fisher’s exact tests for assessment of differences between groups in categorical variables, whenever applicable. We used univariate and multivariate logistic regression models to assess for the association between *M. tuberculosis* lineages and patients’ clinical and demographic characteristics. The associations were assessed for Lineage 2 compared to all other lineages (Lineages 1, 3 and 4), adjusting for age, sex, disease category and HIV status. All statistical analyses were performed in R 3.5.0 [29].

## Results

### Patients’ demographic and clinical characteristics

The patients’ demographic and clinical information in our study included; age, sex, geographical location, HIV and disease category (new or retreatment case). Table 1 describes patients’ characteristics of the study population. The proportions of the observed and missing data for the study population are summarized in S2 Fig.

**Table 1.** Clinical and demographic characteristics of the TB cases

| Characteristics | Valid Proportion % | Total (%)<br>n = 2039 |
| --- | --- | --- |
| **Age, median (IQR)** |  |  |
| 28 (20-38) |  |  |
| **Age groups (years)** |  |  |
| Child age (< 15) | 9.87 | 193 (9.47) |
| Young age (15 - 24) | 29.67 | 580 (28.45) |
| Early adult (25 - 44) | 47.98 | 938 (46.00) |
| Late adult (45 - 64) | 10.03 | 196 (9.61) |
| Old age (> 65) | 2.46 | 48 (2.35) |
| Not available |  | 84 (4.12) |
| *total n = 1955* |  |  |
| **Sex** |  |  |
| Female | 32.40 | 645 (31.63) |
| Male | 67.60 | 1346 (66.01) |
| Not available |  | 48 (2.35) |
| *total n = 1991* |  |  |
| **HIV status** |  |  |
| Negative | 67.71 | 1086 (53.26) |
| Positive | 32.23 | 517 (25.36) |
| Indeterminate | 0.06 | 1 (0.05) |
| Not available |  | 435 (21.33) |
| *total n = 1604* |  |  |
| **Patient category** |  |  |
| New case | 83.95 | 1679 (82.34) |
| Retreatment | 16.05 | 321 (15.74) |
| Not available |  | 39 (1.91) |
| *total n = 2000* |  |  |
| **Geographical zone** |  |  |
| Central | 1.10 | 22 (1.08) |
| Coastal | 51.55 | 1029 (50.47) |
| Lake | 17.94 | 358 (17.56) |
| Northern | 20.19 | 403 (19.76) |
| S. Highlands | 8.12 | 162 (7.95) |
| Western | 0.50 | 10 (0.49) |
| Zanzibar | 0.60 | 12 (0.59) |
| Not available |  | 43 (2.11) |
| *total n = 1996* |  |  |

IQR, interquartile range

Our study population consisted of TB patients ranging between the age of 2 and 82 years with a median age of 28 years (interquartile range [IQR] 20–38). To further probe the age distribution in the study population, we categorized the TB patients into five different age groups (Table 1). We detected approximately three-quarters of the TB cases to occur among the “young age” and “early adult” age groups. Our observation suggests that TB incidence in Tanzania like in other high burden settings [30] is largely contributed by ongoing transmission (rapid progression upon exposure to infection) as opposed to reactivation (longer latency). Further, our findings corroborate with the national TB notification rates in that about 10% of the TB cases are pediatric cases (< 15 years) [31].

Similar to other settings [1], we identified a higher proportion of male TB cases compared to female TB cases. However, the male-to-female ratio in our study population is higher than the national estimates for the two years of sampling (2.2:1 vs., 1.4:1). The striking gender imbalance among TB cases seems to peak at adolescence onwards and is less pronounced among pediatric TB cases (S1 Table). Additionally, a-third (32.2%, 517/1604) of the TB cases with available HIV status were HIV co-infected. In contrast TB/HIV co-infected cases were more likely to be female (44.5%, CI 38.3-50.7% vs., 25.8%, 95% CI 20.6-31.0%) which is consistent with HIV being generally more prevalent in females than males [32]. We found that our study population comprised 16.1% (321/2000) of TB retreatment cases, which was four-fold higher than the overall countrywide notifications [31]. Finally, more than half (51.6%, 1029/1996) of the TB patients in our study population were diagnosed in the Coastal zone of Tanzania and about 40% were either diagnosed in the Lake and Northern zones. In addition to higher TB notification rates, the three former mentioned geographical zones contain the country’s zonal TB reference laboratories. The remaining 10% of the patients were diagnosed in any of the remaining four geographical zones.

### Main MTBC lineages in Tanzania

Using SNP-typing, we detected four of the seven known MTBC lineages (Fig 1), albeit at varying proportions. In our study setting, Lineage 4 and Lineage 3 were the most frequent (866, 42.5% and 681, 33.4%, respectively) followed by Lineage 1 (336, 16.5%). Lineage 2 was the least frequent (92, 4.5%). The remaining 64 clinical isolates (3.1%) could not be assigned into any of the MTBC lineages. Of the seven geographical zones, four (Coastal, Northern, Lake and Southern Highlands) were highly represented with more than 100 clinical strains each (Table 2). The distribution of the *M. tuberculosis* lineages varied within the geographical zones (Fig 1 and S3 Fig). Our findings reveal that Lineage 1 strains were more frequent in the Lake zone compared to the overall average frequency (20.9% vs. 16.8%), whereas the frequency of Lineage 3 in this zone was lower (27.6% vs. 34.3%) compared to other geographical zones. By contrast, Lineage 4 was the most predominant in all geographical zones and showed relatively similar frequencies across the zones.

**Fig 1.**
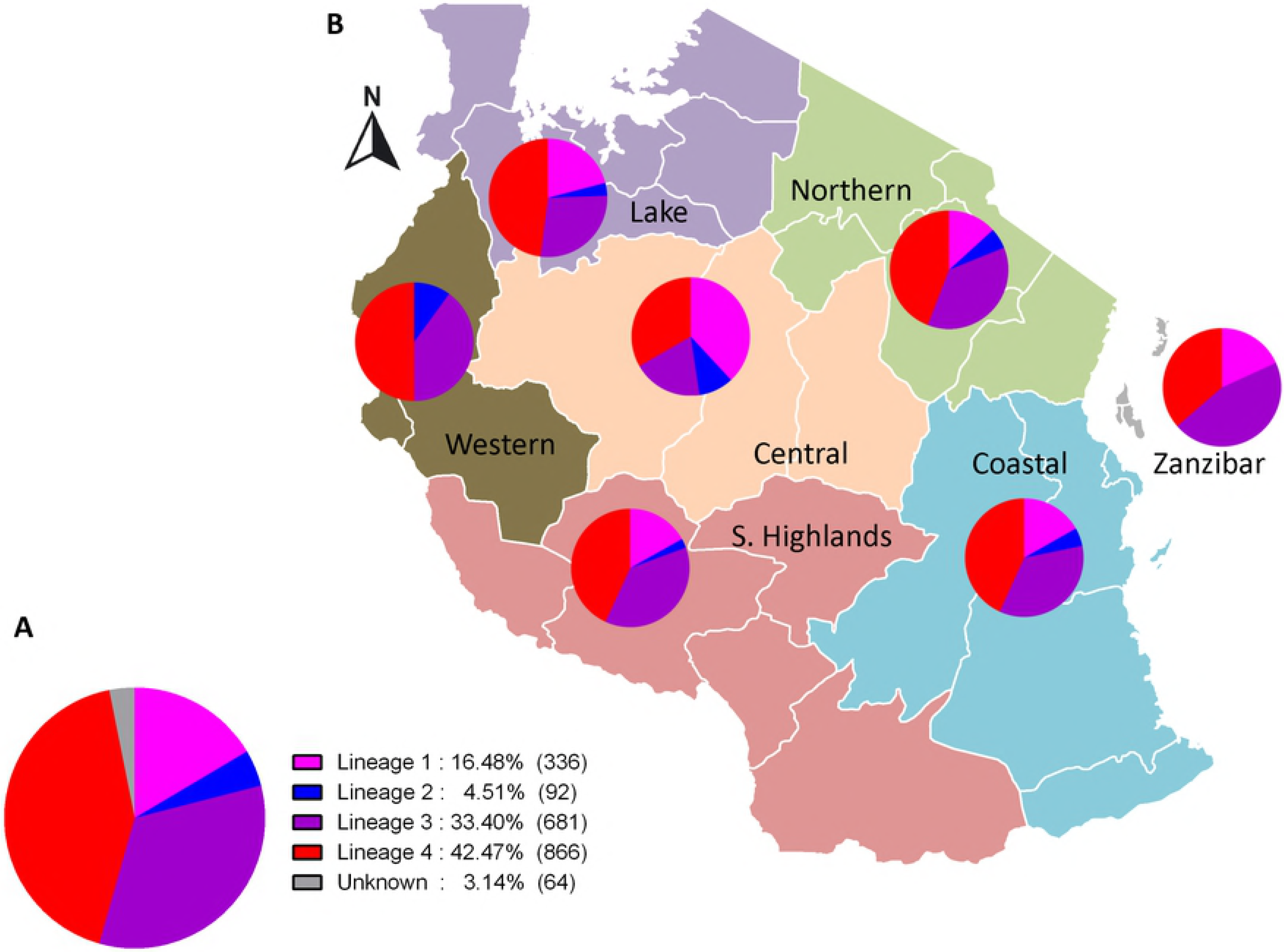
MTBC lineages in Tanzania. A. MTBC lineage classification of 2,039 nationwide clinical strains. B. MTBC lineage frequencies and geographical distribution in Tanzania.

**Table 2.** *M. tuberculosis* lineage distribution across geographical regions in Tanzania

| Geographical Zone | Lineage |  |  |  | Total |
| --- | --- | --- | --- | --- | --- |
|  | L1 (%) | L2 (%) | L3 (%) | L4 (%) |  |
| Central | 8 (38.1 ) | 2 (9.5 ) | 4 (19 ) | 7 (33.3 ) | 21 |
| Coastal | 168 (16.8 ) | 50 (5 ) | 350 (35 ) | 432 (43.2 ) | 1000 |
| Lake | 72 (20.9 ) | 12 (3.5 ) | 95 (27.6 ) | 165 (48 ) | 344 |
| Northern | 52 (13.3 ) | 22 (5.6 ) | 145 (37 ) | 173 (44.1 ) | 392 |
| S. Highlands | 27 (16.9 ) | 4 (2.5 ) | 60 (37.5 ) | 69 (43.1 ) | 160 |
| Western | 0 (0 ) | 1 (10 ) | 4 (40 ) | 5 (50 ) | 10 |
| Zanzibar | 2 (18.2 ) | 0 (0 ) | 5 (45.5 ) | 4 (36.4 ) | 11 |
| **Total** | 329 (17 ) | 91 (4.7 ) | 663 (34.2 ) | 855 (44.1 ) | 1938 |

L1, Lineage 1; L2, Lineage 2; L3, Lineage 3; L4, Lineage 4

### Sublineages

After we detected four main *M. tuberculosis* lineages, we next investigated the respective sublineages within Lineage 1, 3 and 4 using spoligotyping. Lineage 2 strains were excluded from this analysis since the global strains almost exclusively belong to one spoligotype family, Beijing with almost identical fingerprint pattern. We identified 24 spoligotypes (SITs) among the 107 clinical strains analyzed (S2 Table). Twenty six (24.3%) of the strains could not be assigned to any of the existing spoligotypes in the SITVITWEB database and therefore described as orphan spoligotypes. Several spoligotypes were identified within each of the three lineages. Lineage 1 strains mainly belonged to EAI5 spoligotype. On the other hand, CAS1_Kili was the most predominant spoligotype among the Lineage 3 strains. Within Lineage 4 strains, LAM, T, and H families were detected and expectedly the LAM sublineage, particularly LAM_ZWE was the most prevalent.

### Associations between lineages and patients’ characteristics

Having described the circulating main lineages of the *M.tuberculosis* we then assessed the relationship between the lineages and patients’ characteristics (Table 3). We detected a higher proportion of female sex among TB patients infected with Lineage 2 (52.1%) compared to those infected with the other three lineages (range from 31% to 34.5%, p = 0.009). Moreover, we observed that retreatment cases were frequently infected with Lineage 2 strains (26.8%), which was twofold higher compared to Lineage 1 and 4 strains (p < 0.001). We found no evidence for association between lineages and patients’ characteristics such as age and HIV status (Table 3).

Lineage 2 has previously been associated with retreatment cases, drug resistance and lately also with female sex [17,27]. We therefore investigated if similar associations exist in our study population using a subset of TB cases with complete clinical and demographic information (n = 1515). To assess these associations we performed logistic regression analyses comparing Lineage 2 to all other lineages pooled together (Table 4). Our analyses revealed Lineage 2 to be independently associated with female sex (adjusted odds ratio [aOR] 2.25; 95% confidence interval [95% CI] 1.38 – 3.70, p < 0.001) and retreatment cases (aOR 1.78; 95% CI 1.00 – 3.02, p = 0.040). We did not detect any association between the lineages and patients’ age and the HIV status.

**Table 3.**
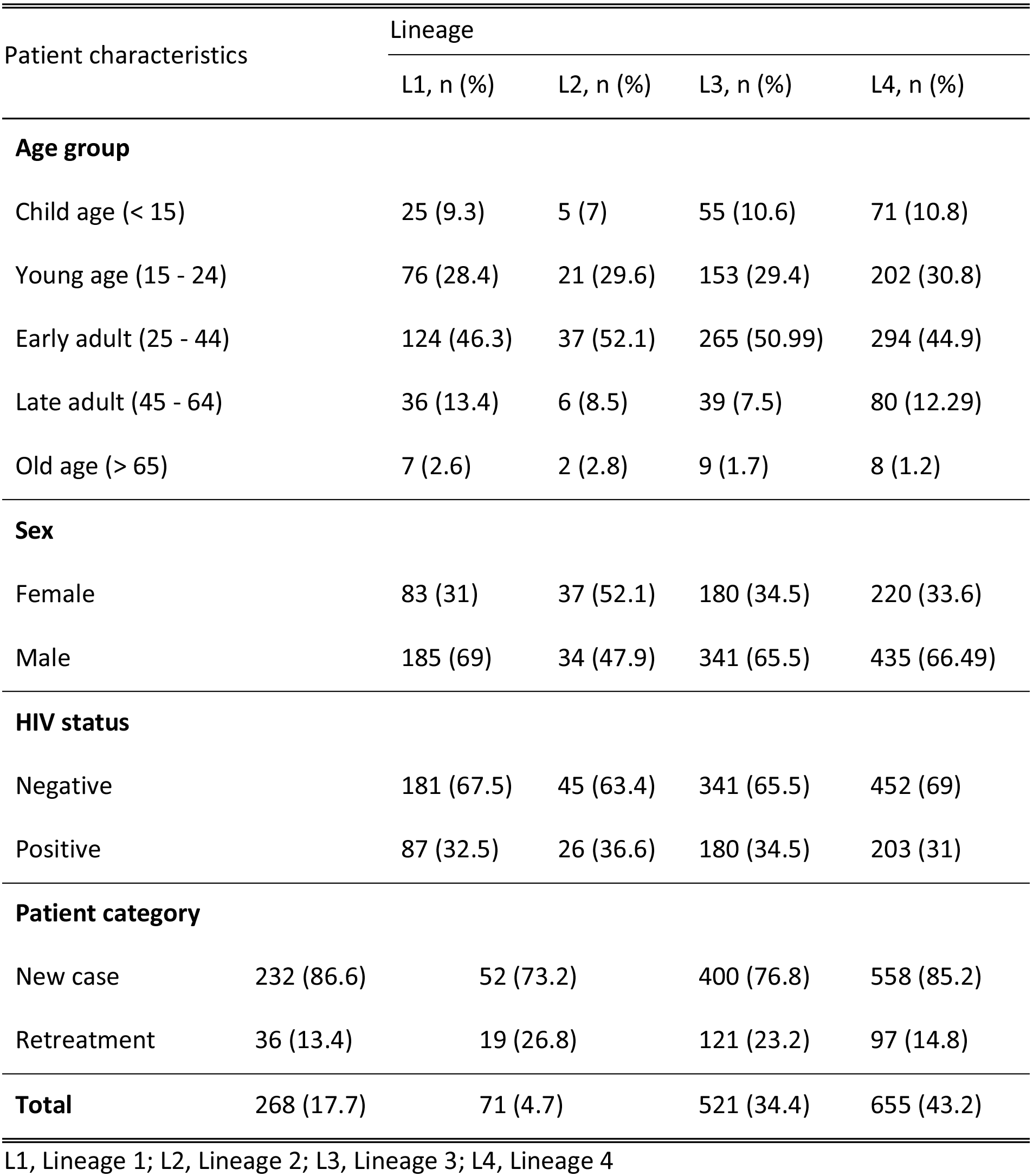
Frequency distribution of *M. tuberculosis* main lineages across patients’ characteristic groups

**Table 4.** Associations of patients’ clinical and demographic characteristics with *M. tuberculosis* Lineage 2 (n = 71) compared to all other lineages (n = 1444)

| Patient characteristics | Lineage 2 | Unadjusted |  | Adjusted |  |
| --- | --- | --- | --- | --- | --- |
|  | n (%) | OR (95% CI) | p value | OR (95% CI) | p value |
| Age, median (IQR) | 27 (20.5 – 38.5) |  |  | 0.99 (0.98 – 1.01) | 0.428 |
| Female sex | 37 (52.1) | 2.17 (1.34–3.51) | 0.002 | 2.25 (1.38 – 3.70) | <0.001 |
| Retreatment case | 19 (26.8) | 1.71 (0.97–2.90) | 0.052 | 1.78 (1.00 – 3.02) | 0.040 |
| HIV positive | 26 (36.6) | 0.98 (0.74 – 1.03) | 0.915 | 1.03 (0.61 – 1.70) | 0.91 |
| Observations |  | 1515 |  | 1515 |  |

IQR, Interquartile range; OR, Odds ratio; 95% CI, 95% confidence interval.

### Mutations within *rpoB* gene in retreatment cases

To investigate whether drug resistance was linked to a particular lineage, we included in total 145 retreatment cases for drug resistance genotyping of the *rpoB* gene that confers resistance to rifampicin. Out of these, 112 (77.2%) had no mutations compared to the H37Rv reference gene and 16 (11%) contained at least one mutation, either synonymous (3/16) or non-synonymous (13/16) (S4 Fig and S3 Table). We could not determine mutation status in the *rpoB* gene of 17 (11.7%) retreatment cases due to PCR and sequencing failure. Among the 13 strains detected with non-synonymous *rpoB* mutations, five belonged to Lineage 2, four to Lineage 4, three to Lineage 3 and one was unclassified (S4 Table). Table 4 summarizes the non-synonymous *rpoB* mutations detected.

**Table 4.** Detected mutations on the *rpoB* gene among the retreatment cases

| rpoB mutation | Amino acid change | Comment |
| --- | --- | --- |
| A1198G | T400A | reported |
| A1304T | D435V | reported |
| A1334T | H445L | reported |
| C1333G | H445D | reported |
| C1333T | H445Y | reported |
| T1289C | L430P | reported |
| A1442G | E481A | reported |
| C1294G | Q432E | reported |
| C1349T | S450L | reported |

## Discussion

In this study, we classified the countrywide collection of 2,039 *M. tuberculosis* isolates from smear positive new and retreatment TB cases diagnosed between 2012 and 2013 in Tanzania. Our findings show that the *M. tuberculosis* strains in Tanzania are diverse, comprising four main phylogenetic lineages (Lineage 1–4) which occur throughout the country. Specifically, we found that Lineage 4 was the most frequent nationwide, followed by Lineage 3 and 1. Despite Lineage 2’s recent global dissemination [15], it was the least frequent in our study setting. Finally, our analysis on the relationship between *M. tuberculosis* lineages and patients’ characteristics revealed associations of Lineage 2 with female sex and retreatment TB cases.

Among the 7 human-adapted MTBC lineages, Lineage 4 is the most broadly distributed and occurs at high frequencies in Europe, the Americas and Africa [26,33]. We observe that TB epidemics in Tanzania are also predominated by Lineage 4, which is regarded as the most successful of MTBC lineages [33]. In general, the wide geographical range of Lineage 4 is postulated to be driven by a combination of its enhanced virulence, high rates of human migration linked to its spread and ultimately its ability to infect different human population backgrounds [33,34]. In contrast, Lineage 1 and 3 are known to be mainly confined to the rim of the Indian Ocean [9], which is consistent with our observation that nearly 50% of the *M. tuberculosis* strains in Tanzania belong to these two lineages. This high prevalence of Lineage 1 and 3 likely reflects the long-term migrations between Eastern Africa and the Indian subcontinent [35]. In addition, the distribution and the frequency of Lineage 1 and 3 in the mainland away from the coast of Tanzania did not vary, suggesting spread via internal migrations. Lineage 1 was proposed to have evolved in East Africa prior disseminating out of the continent [12]. Based on this, one might expect higher frequencies of Lineage 1 in the region. Instead, the so called “modern” (TbD1−) lineages (4 and 3 in this case) dominate in Tanzania despite presumably being introduced into the African continent only post-contact [33,36]. This illustrates the ability of “modern” lineages to thrive in co-existence with the pre-existing “ancient” (TbD1+) lineages such as Lineage 1 in our case, perhaps because of the comparably higher virulence [16,37]. The neighboring countries of Tanzania on the other hand show comparable *M. tuberculosis* lineage composition [38,39], indicating common demographic histories and ongoing exchanges that resulted into distinct *M. tuberculosis* populations. The frequency of Lineage 2–Beijing in Tanzania, like in most parts of the continent except for South Africa [38,39] is relatively low, despite the long-standing African-Asian contacts [39]. Evidence from recent studies show that Lineage 2-Beijing was only recently introduced into Africa [15,40].

The burden of TB disease is generally higher in males [1,41], rendering male sex as a potential risk factor for TB. Furthermore, the male bias among TB patients is also observed in settings with no obvious sex-based differences in health-seeking behavior [42]. Whilst we show similar trends in this study setting overall, our findings reveal that the proportion of females was higher among TB patients infected with Lineage 2. This finding is consistent with several other previous studies conducted in different settings [17,27,43]. Social and physiological factors predisposing males to higher risk of TB have been indicated [44]. On the one hand, these include risk behaviors such as substance abuse (alcoholism, tobacco smoking) and gender specific roles such as risk occupations (e.g., mining) that are male dominated and known to increase the risk for TB. On the other hand, genetic makeup and sex hormones might contribute to the differences in TB susceptibility among females and males, as epidemiological and experimental studies have suggested female sex hormones to be protective [44]. These observations would propose that the sex imbalance in TB to emerge after the onset of puberty. Of note, we observe less sex imbalance in “child” age group (<15 years) which also corroborates the national notification rates [31]. However, this observation can be confounded by BCG vaccination which might be most effective in this age group. Despite the high prevalence of HIV among young females in sub-Saharan Africa [32] and HIV being the strongest risk factor for TB, TB burden remains higher in males. While social and physiological aspects play an important role, findings from this study and others previously conducted in Nepal and Vietnam [17,27] suggest that bacterial factors could disrupt the trends towards male bias in TB, a finding which warrants further investigation. Our hypothesis is that because of higher virulence, Lineage 2 strains are able to overcome the resistance poised by female sex which could explain the less pronounced sex imbalanced.

In addition to its association with female sex, Lineage 2 was also associated with retreatment TB cases [43]. A retreatment case in our study population represented recurrent TB case either due to relapse or reinfection. We hypothesized that this observation was possibly linked to drug resistance, given the previous reported association between Lineage 2 and drug resistance [45]. However, we detected only 9% (13/145) of strains among the retreatment subset tested to contain mutations conferring resistance to rifampicin, five of which belonged to Lineage 2. These findings would suggest that retreatment cases are mainly driven by reinfection as opposed to treatment failure or relapse.

Finally, as evidenced by the age distribution of TB cases in our study setting, recent or ongoing transmission in high burden countries is the main contributor to the TB burden rather than disease reactivation [30]. Additionally, an association with young age has been employed as an epidemiological proxy for highly transmissible strains and faster rates of disease progression [46,47]. In this study, we did not detect any differences in median age of TB patients infected with different lineages (S5 Fig), an observation that could speak for high ongoing transmission rates in general, irrespective of lineage.

Our study is limited by focusing on a convenient collection of archived *M. tuberculosis* clinical isolates (representing 1.6% of all TB cases in 2012 and 2013) sampled from TB cases as part of countrywide drug resistance surveillance. Therefore, the strength or lack of associations between lineages and patients’ characteristics could likely be affected by the sampling. In addition, most of the geographical zones were underrepresented which could in turn underestimate the respective regional lineage composition and the overall countrywide distribution. Systematic sampling would allow for better resolution on the distribution patterns, the frequencies and on epidemiological consequences of *M. tuberculosis* lineages, which might partially determine the regional specific epidemics.

In conclusion, this study addresses the countrywide *M. tuberculosis* population structure based on SNP-typing. We show that *M. tuberculosis* population in Tanzania is diverse with four of the seven known lineages detected. This study sets the stage for further in depth investigations on epidemiological impact of *M. tuberculosis* lineages in Tanzania.

## Acknowledgements

We would like to thank the National Tuberculosis Leprosy Programme (NTLP) through the Central Reference Laboratory (CTRL) for permission to use the *M. tuberculosis* isolate collection for this study.

## Supporting Information

**S1 Fig. Study population flowchart.**

**S2 Fig. Patients’ data included in the study.** Proportion of observed and missing data for the variables included in the study

**S3 Fig. MTBC lineage proportions.** Distribution of *M. tuberculosis* lineages across different regions of Tanzania. Size of the circle is proportional to the number of isolates analyzed from the regions.

**S4 Fig. Flowchart of genotyped strains for *rpoB* mutations.** A subset of *M. tuberculosis* strains from retreatment cases included for *rpoB* drug resistance genotyping

**S5 Fig. Patients’ age distribution across MTBC lineages.** The age distributions of TB patients grouped by infecting MTBC lineage.

**S1 Table. Sex distribution across different age groups of TB patients.**

**S2 Table. Spoligotype patterns of a subset of *M. tuberculosis* clinical strains**

**S3 Table. Mutations detected in the *rpoB* gene**

**S4 Table. Distribution of *rpoB* mutations across the four MTBC lineagesFigurer**

